# Multiomics analysis integrating pyroptosis-related signatures for building a prognostic prediction model in hepatocellular carcinoma

**DOI:** 10.1101/2022.01.24.477487

**Authors:** Jiahao Huang, Yan Li, Ming Chu, Yuedan Wang

## Abstract

Hepatocellular carcinoma (HCC) is a major cause of cancer-related death worldwide and has a poor prognosis. Pyroptosis, which is programmed cell necrosis mediated by gasdermin, participates in the progression of tumors. Recently, multiple omics analysis has been applied frequently to provide comprehensive and more precise conclusions. However, multiomics analysis combining pyroptosis-related signatures in HCC and their correlations with prognosis remain unclear. Here, we identified 42 pyroptosis genes that were differentially expressed between HCC and normal hepatocellular tissues. According to these differentially expressed genes (DEGs), all HCC cases could be divided into two heterogeneous subtypes. Then, we evaluated the prognostic value of differential pyroptosis-related genes to construct a multigene model using The Cancer Genome Atlas (TCGA) cohort. A 22-gene model was built and classified HCC patients in the TCGA cohort into low-risk and high-risk groups by the least absolute shrinkage and selection operator (LASSO) Cox regression method. HCC patients belonging to the low-risk group had significantly higher survival possibilities than those belonging to the high-risk group (p<0.001). Furthermore, the related genes and two groups were analyzed with multiple omics in different molecular layers. The pyroptosis-related gene model was validated with HCC patients from the Gene Expression Omnibus (GEO) cohort, and the low-risk group in GEO showed increased overall survival (OS) time (P=0.018). The risk score was an independent factor for predicting the OS of HCC patients. In conclusion, pyroptosis-related genes in HCC are correlated with tumor immunity and could be used to predict the prognosis of HCC patients.

## Introduction

Hepatocellular carcinoma (HCC) results in >80% of primary liver cancers in the world. HCC also causes a heavy disease burden and is estimated to be the fourth most common cause of cancer-related death worldwide[1]. The main risk factors for HCC are hepatitis B virus and hepatitis C virus infection[2]. Moreover, nonalcoholic steatohepatitis associated with metabolic syndrome is becoming a more frequent risk factor[3]. Currently, there are various treatment options, including surgical resection, chemotherapeutics, immunotherapies, new methods for the delivery of drugs and the use of combination therapy[4]. However, the adjusted incidence rates and death rates have continued to increase[5]. Noninvasive diagnosis is currently challenged by the requirement of molecular information that requires tissue or liquid biopsies[3]. Thus, it is still urgent to find ideal combination therapies or advanced detection methods for early-stage hepatocellular carcinoma.

Multiomics analysis provides an integrative analysis to maximize comprehensive biological insight across molecular layers[6]. A novel form of cell-regulated necrosis, pyroptosis, which is mainly induced by gasdermin, plays a crucial role in cancer and hereditary diseases[7]. Pyroptosis is an inflammatory form of cell death that is characterized by cellular swelling and bubble-like protrusions[8]. Pyroptosis can be triggered by canonical caspase-1 inflammasomes or by the activation of caspase-4, -5 and -11 by cytosolic lipopolysaccharide[9]. Then, canonical caspase-1 inflammasomes cleave the effector molecule gasdermin D (GSDMD) and promote its oligomerization to form large pores in the plasma membrane, causing cell death[10]. In addition, some cytokines, such as IL-18 and IL-1β, were active during pyroptosis. Moreover, pyroptosis has a crucial role in the proliferation and migration of cancer regulated by molecules such as noncoding RNAs[11]. Furthermore, it has been revealed that pyroptosis-induced inflammation triggers robust antitumor immunity and can synergize with checkpoint blockade[12]. These findings collectively demonstrate that pyroptosis has significant roles in the development and antitumor processes. However, its specific functions with multiple omics in HCC have not been reported. Thus, we performed a multiomics study to determine the functions of pyroptosis-related genes in HCC, explore the gene copy number variants, mutations, immunocyte correlations, tumor stem cell correlations and drug sensitivities of related genes and two different risk groups, and establish a robust prognostic model based on pyroptosis for the detection of early-stage HCC.

## Materials and Methods

### Dataset Collection

The HCC RNA-seq count and clinical profiles were obtained from the TCGA GDC database (https://portal.gdc.cancer.gov/) and the GEO database (https://www.ncbi.nlm.nih.gov/geo/, ID: GSE20140). The FPKM data normalized from the RNA-seq count were transformed to log2(TPM+1) for further analysis. The data of cope number variation and simple nucleotide variation were downloaded from TCGA GDC database (https://portal.gdc.cancer.gov/). All expression data were normalized before analysis. Patients were excluded if they died within 30 days or did not have prognostic information.

### Analysis of differential pyroptosis-related genes in HCC

Fifty-five pyroptosis-related genes were collected from prior articles. The differentially expressed genes (DEGs) in HCC samples and matched normal tissues were analyzed by the “limma” package in R software and visualized by a heatmap with an adjusted p value < 0.05. The protein–protein interaction network for DEGs was generated using the STRING website (https://cn.string-db.org/). Gene Ontology (GO) and Kyoto Encyclopedia of Genes and Genomes (KEGG) pathway enrichment analyses were applied to explore the molecular mechanisms of risk using the R “clusterProfiler” package.

### Construction and validation of the prediction model for HCC

Cox regression analysis was used to screen the prognostic DEGs. The screened genes were further narrowed down by the LASSO Cox regression model (R “glmnet” package) to develop the prognostic model. After standardization and normalization of the expression data, the risk score formula was calculated based on a screened 22-gene signature as follows: Σ*^7_i_^ Xi* × *Yi* (*X*: coefficients, *Y*: gene expression level). Subsequently, the patient samples from the TCGA and GEO cohorts were divided into low- and high-risk groups based on the model. Kaplan–Meier survival curves were depicted to predict the clinical outcomes in the two groups by the R “survival” package. The R “survminer” and “timeROC” packages were applied to assess the survival and prognosis of patients.

### Multi-Omics Data Analysis

To determine the mutation of DEGs in samples, the R “maftools” package was applied. The R “CIBERSORT” package was used to assess immune cell infiltration in the samples. Then, the R “ggplot2” was used to visualize the correlation of immune cells and risk genes. To explore the TME, the scores of immune-related projects were evaluated by the R “estimate” package. The drug sensitivity analysis was performed by the R “pRRophetic” package. The R “ggpubr” and “ggExtra” packages were employed to assay the correlation of the tumor stem cell index and risk.

### Statistical Analysis

One-way ANOVA was applied to calculate the differences in gene expression. The Pearson chi-square test was used to compare the profiles between the two subgroups. Differences in OS between the two subgroups were analyzed by the Kaplan–Meier method with a two-sided long-rank test. Hazard ratios (HRs) were calculated by univariate and multiple Cox regression analyses. All statistical significance was considered as a *p* value less than 0.05. All statistical analyses were performed using R software 4.0.1.

## Results

### Landscape of pyroptosis genes in HCC

Fifty-five pyroptosis-related genes were collected and compared in The Cancer Genome Atlas (TCGA) data from 50 normal and 374 tumor tissues. Together, we identified 42 differentially expressed genes (DEGs) (all *p* <0.05). Among them, 9 genes were downregulated, while 32 other genes were upregulated in the tumor group compared to the normal group. The RNA levels of these genes are presented as heatmaps in Fig. 1A. To further explore the interactions of these differentially expressed pyroptosis-related genes, we conducted a protein–protein interaction (PPI) analysis, and the results are shown in Fig. 1B. The minimum required interaction acore for the PPI analysis was set at 0.9. The correction network containing all pyroptosis-related genes is presented in Fig. 1C. Furthermore, the tumor mutational burden (TMB) of these genes in HCC samples was assayed (Fig. 2A). The frequency of copy number variations (CNVs) of pyroptosis genes in HCC was further evaluated (Fig. 2B).

**Fig. 1:**
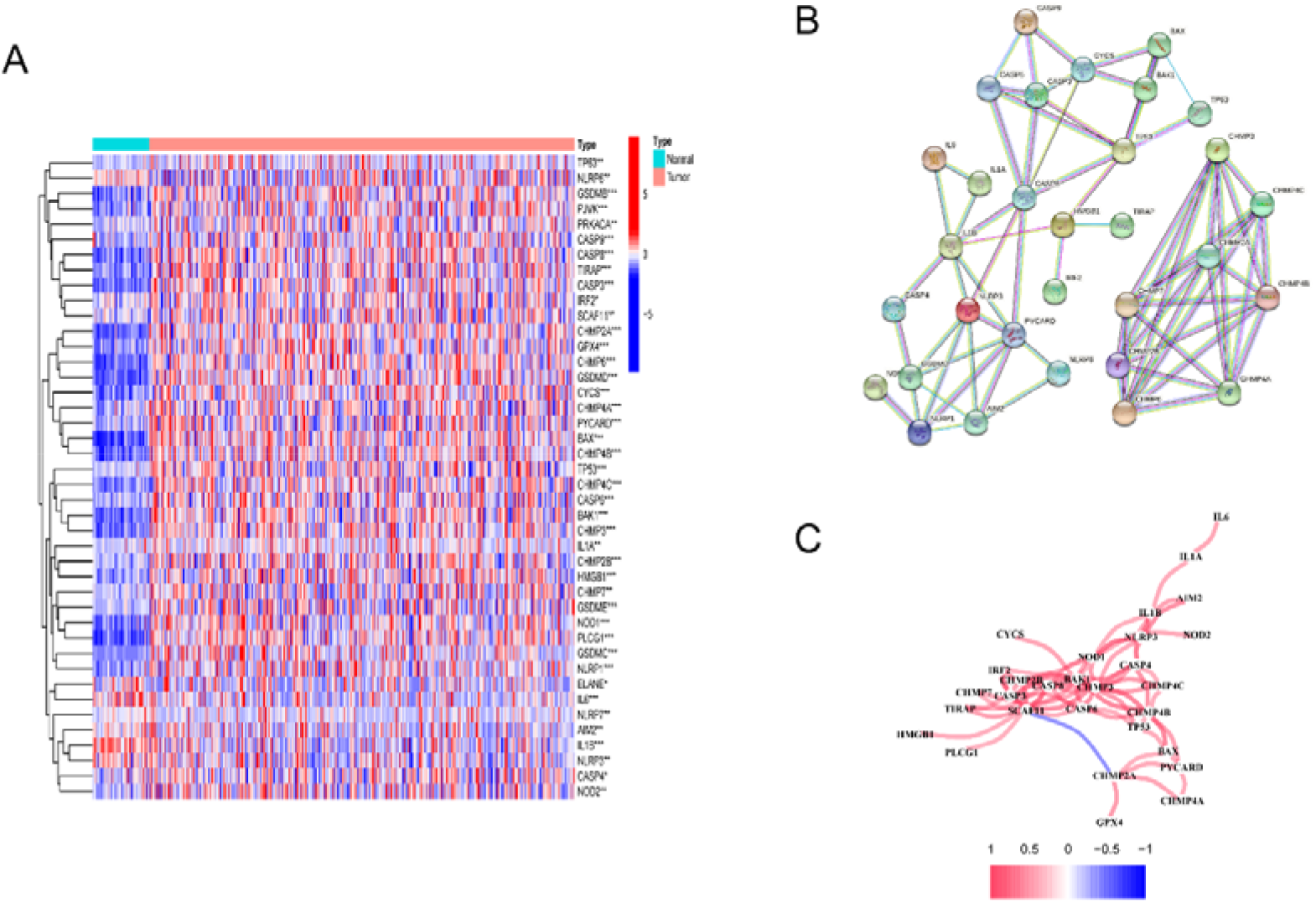
The landscape of pyroptosis-related genes in HCC. **(A)** Heatmap of the differential gene expression between the normal and tumor tissues. **(B)** PPI network of the DEGs. **(C)** The correlation network (red line: positive correlation; blue line: negative correlation. The depth of the colors reflects the strength of the relevance.

**Fig. 2:**
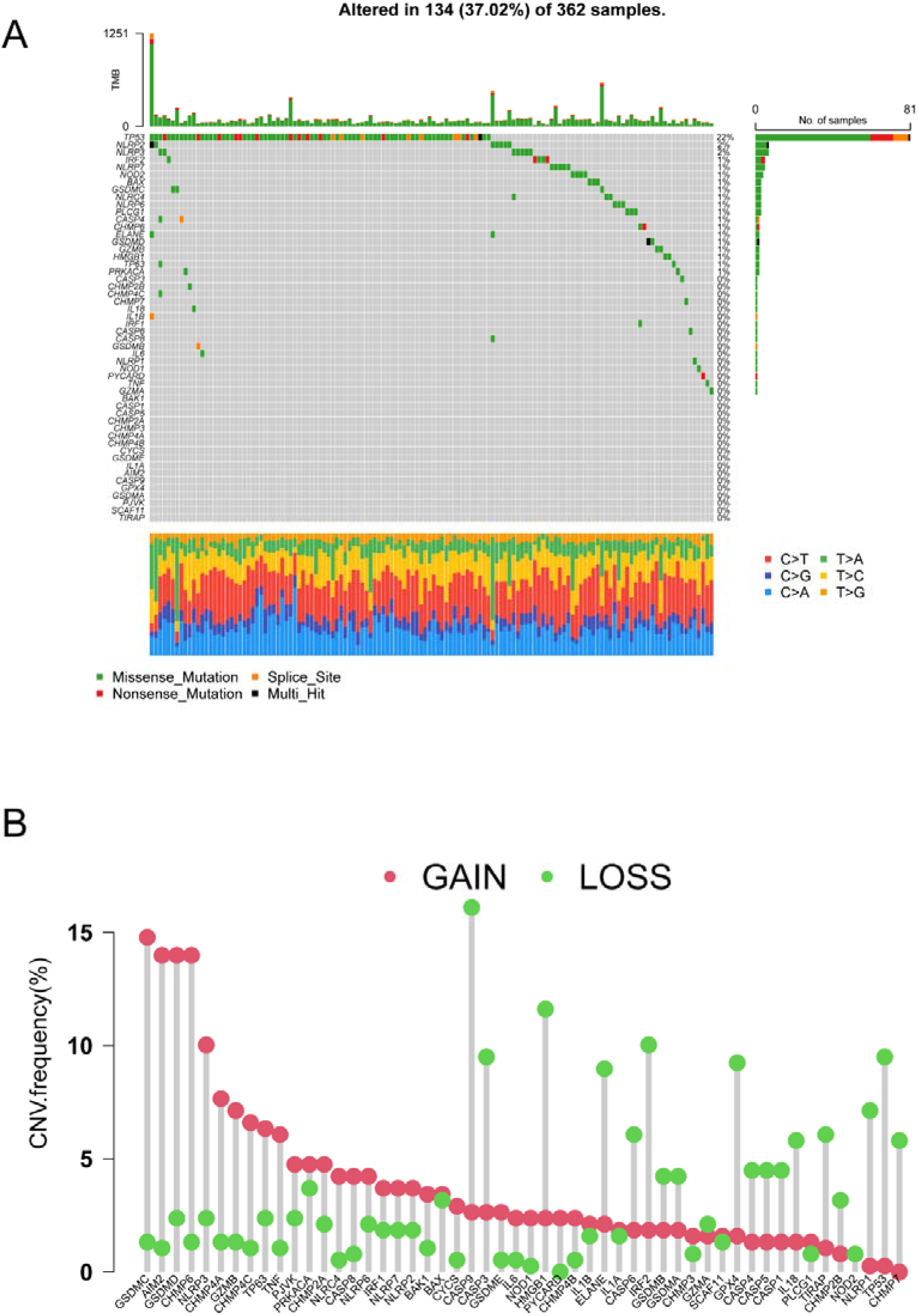
The landscape of DEGs in HCC. **(A)** The simple nucleotide variation of DEGs in HCC samples. **(B)** The cope number variation of DEGs in HCC samples.

### Tumor classification based on the expression level of pyroptosis genes

After removing the normal hepatocellular tissues, we used unsupervised clustering methods to classify the tumor samples into different molecular subgroups based on pyroptosis-related genes. By increasing the clustering variable (*K*) from 2 to 9, we found that when *K*=2, the intragroup correlations were low, indicating that the HCC patients could be well divided into two clusters, termed C1 and C2, based on the 42 DEGs (Fig. 3A, B). Combining the matched clinical profiles, we found that the overall survival (OS) time of the two clusters was significantly different (Fig. 3C). To evaluate the clinical significance of subtypes, clinical outcomes and clinicopathological features were compared between the two clusters, the results showed that the grade of disease was significantly different between the two clusters (*p*<0.001) (Fig. 3D).

**Fig. 3:**
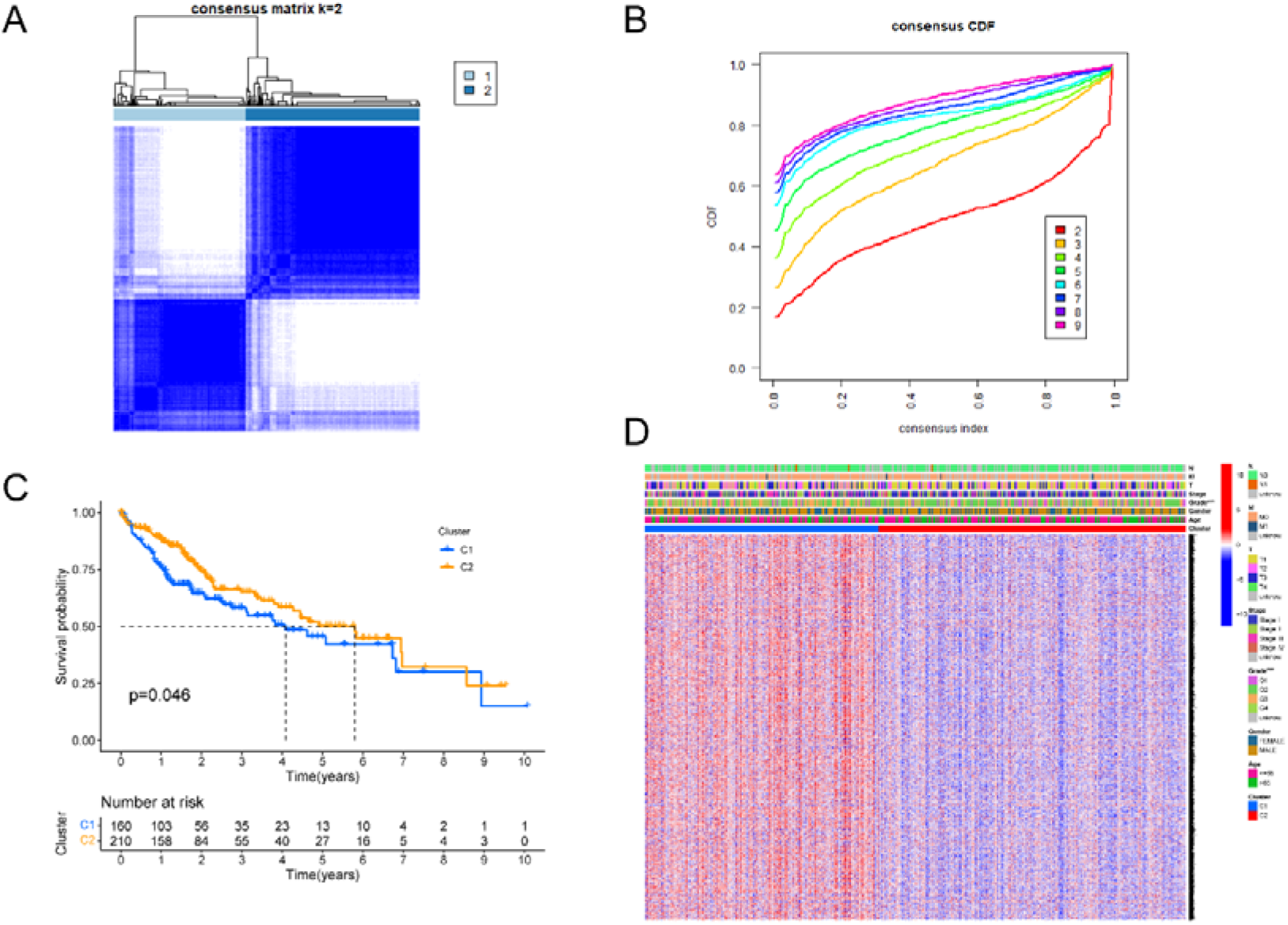
Identification of heterogenetic clusters based on the DEGs. **(A)** Consensus cluster matrix of HCC samples when k = 2. **(B)** The cumulative distribution function curves with k from 2 to 9. **(C)** Kaplan–Meier analysis for overall survival of the two clusters. **(D)** The clinicopathologic differences of the two clusters.

### Establishment of an accurate prognostic model using the TCGA cohort

The differentially expressed genes were screened between the two clusters. In addition, 79 HCC samples from the Gene Expression Omnibus (GEO) cohort (GSE20140) were selected. Then, we intersected the genes of the TCGA cohort, GEO cohort and differential genes from two clusters. Furthermore, the intersected genes of the TCGA cohort combining matched clinical information were analyzed with univariate Cox regression analysis to screen for survival-related genes. The *p* value filter was 0.001. The majority of survival-related genes were associated with increased risk with HRs>1 (Table 1). Through least absolute shrinkage and selection operator (LASSO) Cox regression analysis, 22 genes were selected to be modeled according to the optimum λ value (Fig. 4A, B). The risk forecasting formula was calculated as follows: = (−0.004\**SPP1* exp.) + (0.143\**MYCN* exp.) + (−0.03\**PON1* exp.) … + (0.11\**MT3* exp.) (Table 2). Based on the median score calculated by the risk score formula, 374 HCC patients from TCGA cohort were trained into low- and high-risk subgroups (Fig. 4C). Principal component analysis (PCA) showed that patients with different risks were well separated into two clusters (Fig. 4D). There was a significant difference in OS time between the low- and high-risk groups (P<0.001) (Fig. 4E). The time-dependent receiver operating characteristic (ROC) analysis showed that the areas under the ROC curve (AUCs) were generally higher than 0.8 at 1, 3, and 5 years, demonstrating that the prognostic model has high accuracy and sensitivity (Fig. 4F).

**Table.1.** The survival-related genes.

**Table.2.** The risk forecasting formula.

**Fig. 4:**
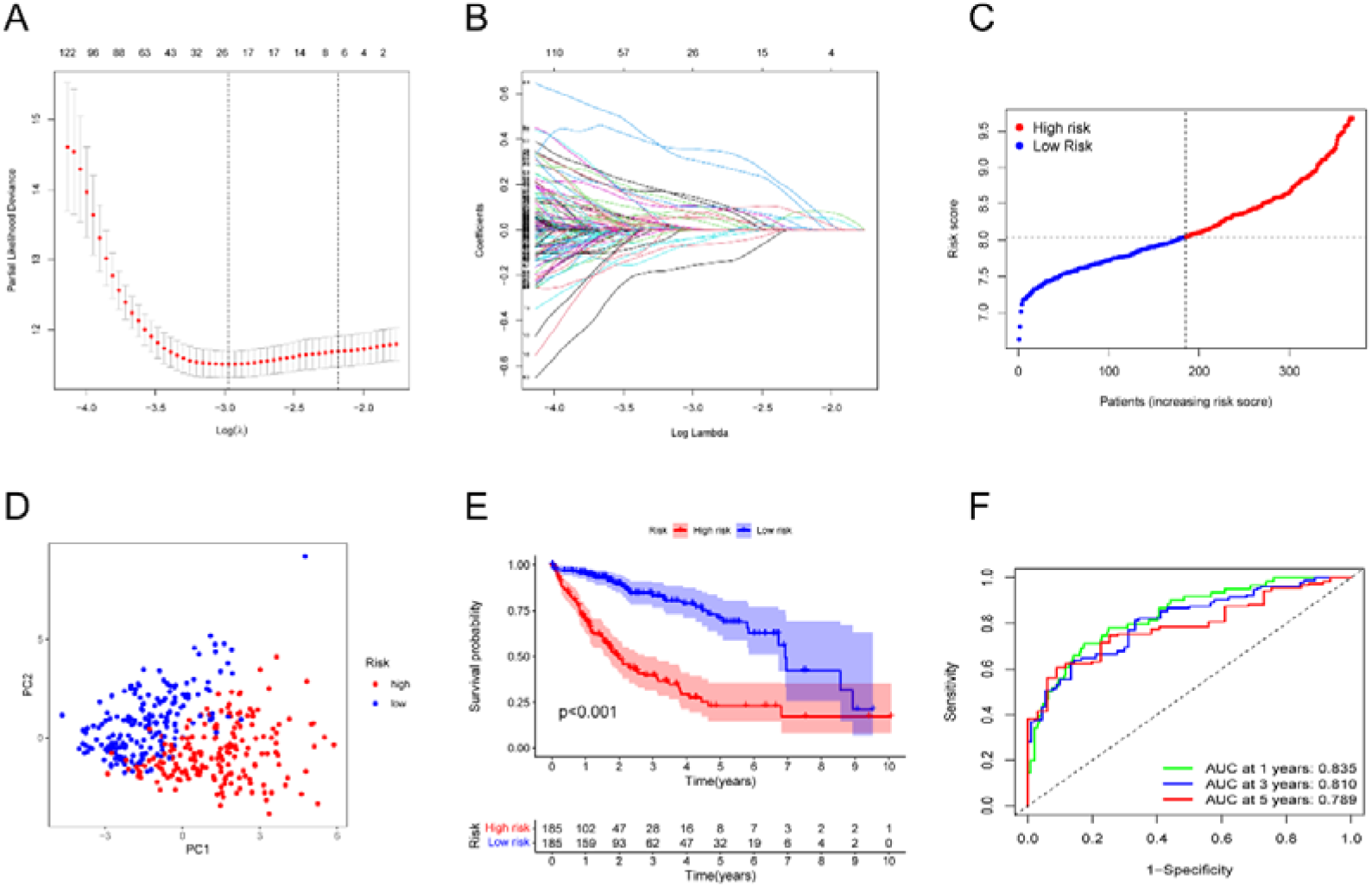
Construction of the prognostic model. **(A)** The optimal parameter (λ) was chosen by cross-validation. **(B)** LASSO coefficient plot of the prognostic genes. **(C)** Risk score analysis in HCC patients. **(D)** PCA plot for HCC patients based on the risk score. **(E)** KM analysis of overall survival in the two groups. **(F)** ROC analysis to evaluate the predictive efficiency.

### Internal and external validation of the prognostic model using the GEO cohort

HCC patients from a Gene Expression Omnibus (GEO) cohort (GSE20140) were utilized as the validation set. The gene expression data were normalized before analysis. According to the median risk score from the TCGA cohort, patients from the GEO cohort were also divided into low- and high-risk groups (Fig. 5A). The PCA showed satisfactory separation between the two subgroups (Fig. 5B). Similar to the TCGA cohort, Kaplan–Meier analysis indicated that the low-risk group had better overall survival than the high-risk group (*p*=0.018) (Fig. 5C). Moreover, the ROC curve analysis showed that the AUC was 0.846 for 5 years, 0.839 for 7 years, and 0.817 for 9 years (Fig. 5D).

**Fig. 5:**
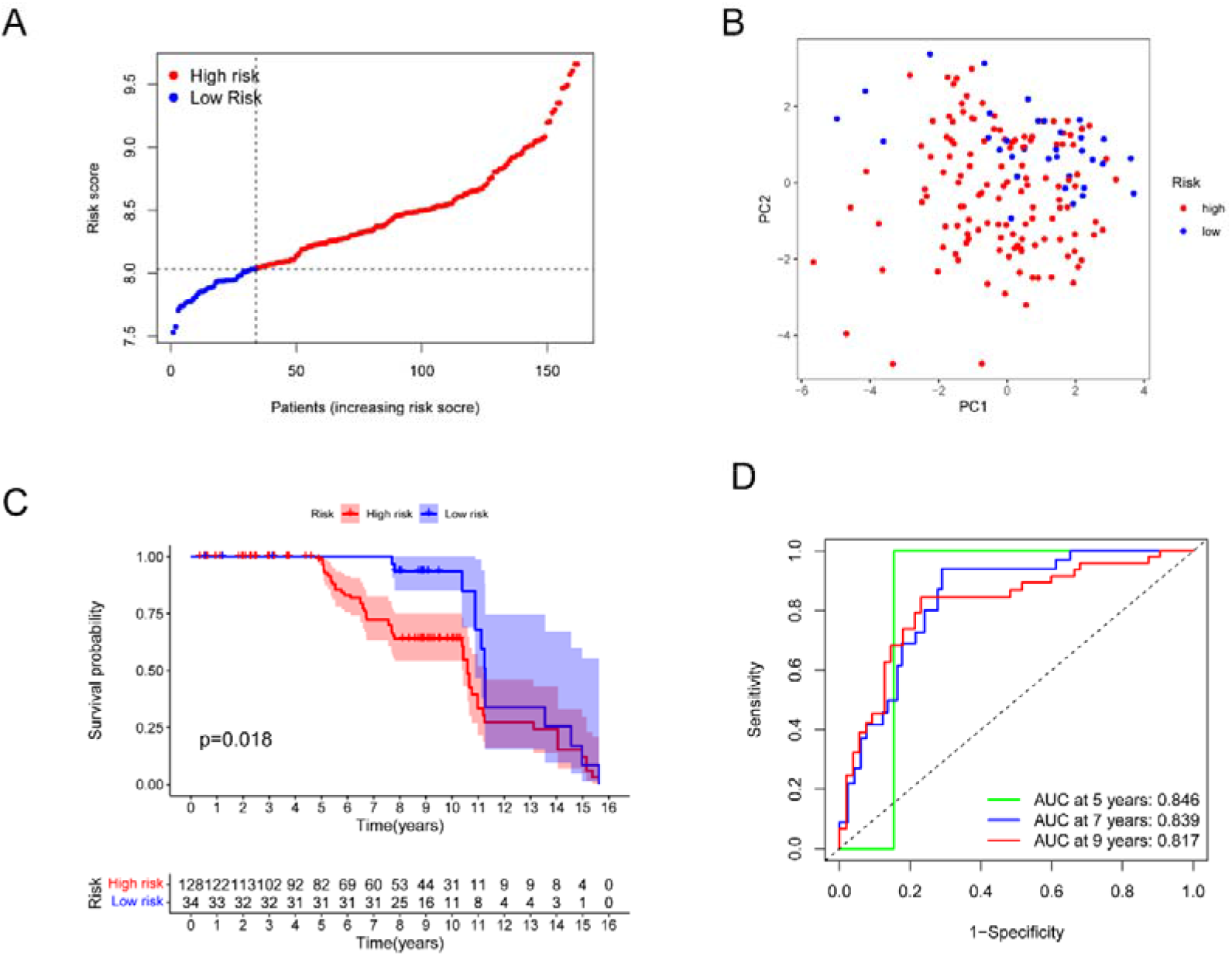
Validation of the prognostic model in the GEO cohort. The figure legends are the same as those in Fig. 4.

### The prognostic value of the established risk model

First, the model genes were analyzed by univariate Cox regression to evaluate the prognostic value of some features, such as risk score, age, and disease stage. The results showed that the risk score and disease stage were significant prognostic factors for patients (*p*<0.05) (Fig. 6A). Subsequently, we included the two factors in further multivariate analysis, which indicated that the risk score can serve as an independent prognostic factor (*p*<0.05) (Fig. 6B). Moreover, we generated a risk heatmap of clinical characteristics based on the model genes. The map showed that the T staging, stage and grade of HCC were distributed diversely between the high- and low-risk groups divided by these model genes (p<0.001) (Fig. 6C).

**Fig. 6:**
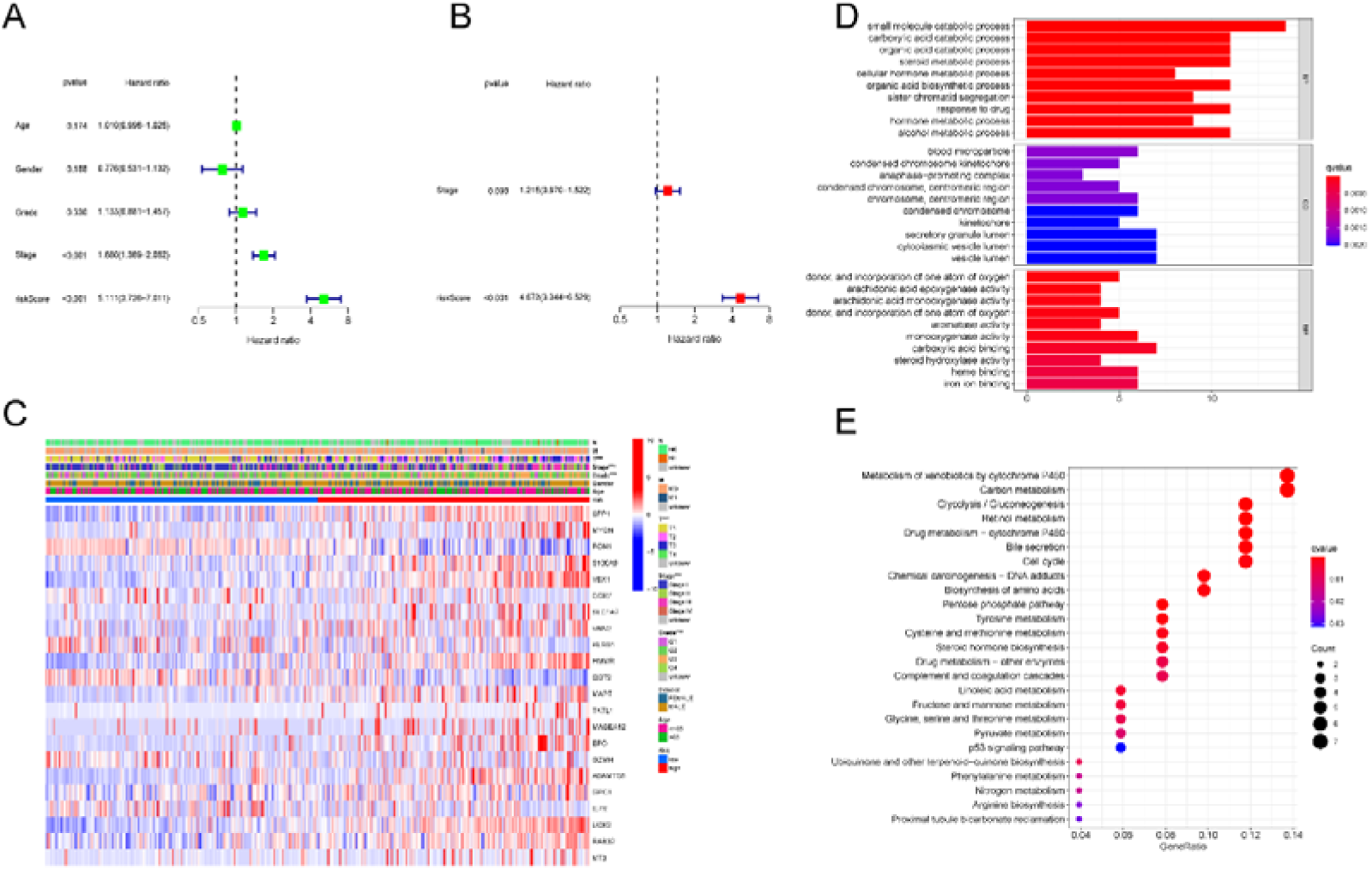
The value of prognosis and the enrichment analysis. **(A, B)** Univariate and multivariate Cox analyses based on the risk model. **(C)** Heatmap for the correlations between clinical profiles and risk groups (\*\*\**p*<0.001). **(D, E)** GO and KEGG pathway enrichment analyses of the DEGs.

### Identification of DEGs and functional analysis

The differential gene expression (DEGs) was analyzed between the two risk subgroups defined by the risk model. The classical method “limma” R package was applied to identify the DEGs. The filter standard was set at |log2FC | ≥ 1 and the adjusted *p* value (FDR) < 0.05. Together, there were 73 DEGs between the two subgroups (Table 3). Subsequently, these 73 DEGs were further enriched by Gene Ontology (GO) and Kyoto Encyclopedia of Genes and Genomes (KEGG) pathway analyses. The enriched results indicated that biological functions contributing to the risk difference were mainly related to material metabolism and molecule secretion (Fig. 6D, E).

**Table.3.** The DEGs between the two subgroups.

### Comprehensive biological analysis with multiple dimensions

The immune cell infiltration in HCC samples from the TCGA cohort was evaluated, and the correlation of model genes and the infiltrated immune cells was further analyzed. The results indicated that CD8 T cells and activated memory CD4 T cells had a strong positive correlation with the GZMH gene, while resting memory CD4 T cells had a strong negative correlation with *GZMH* (Fig. 7A). Moreover, we evaluated the tumor microenvironment (TME) of HCC samples according to immune-related scores (including stromal score, immune score and estimate score). The difference between the scores of the two risk subgroups was further compared, which showed that the low-risk group had higher scores in all three immune-related dimensions than the high-risk group (Fig. 7B). Furthermore, the risk score generated by the risk model had a significant positive correlation with the tumor stem cell index (RNAss) (Fig. 7C). Finally, the sensitivity of some drugs with therapeutic potentialities was investigated in the two risk groups. It was found that some drugs, such as nilotinib, bortezomib and dasatinib, had significantly different IC50 values between the low- and high-risk groups, and the high-risk group had a lower IC50 value (Fig. 8).

**Fig. 7:**
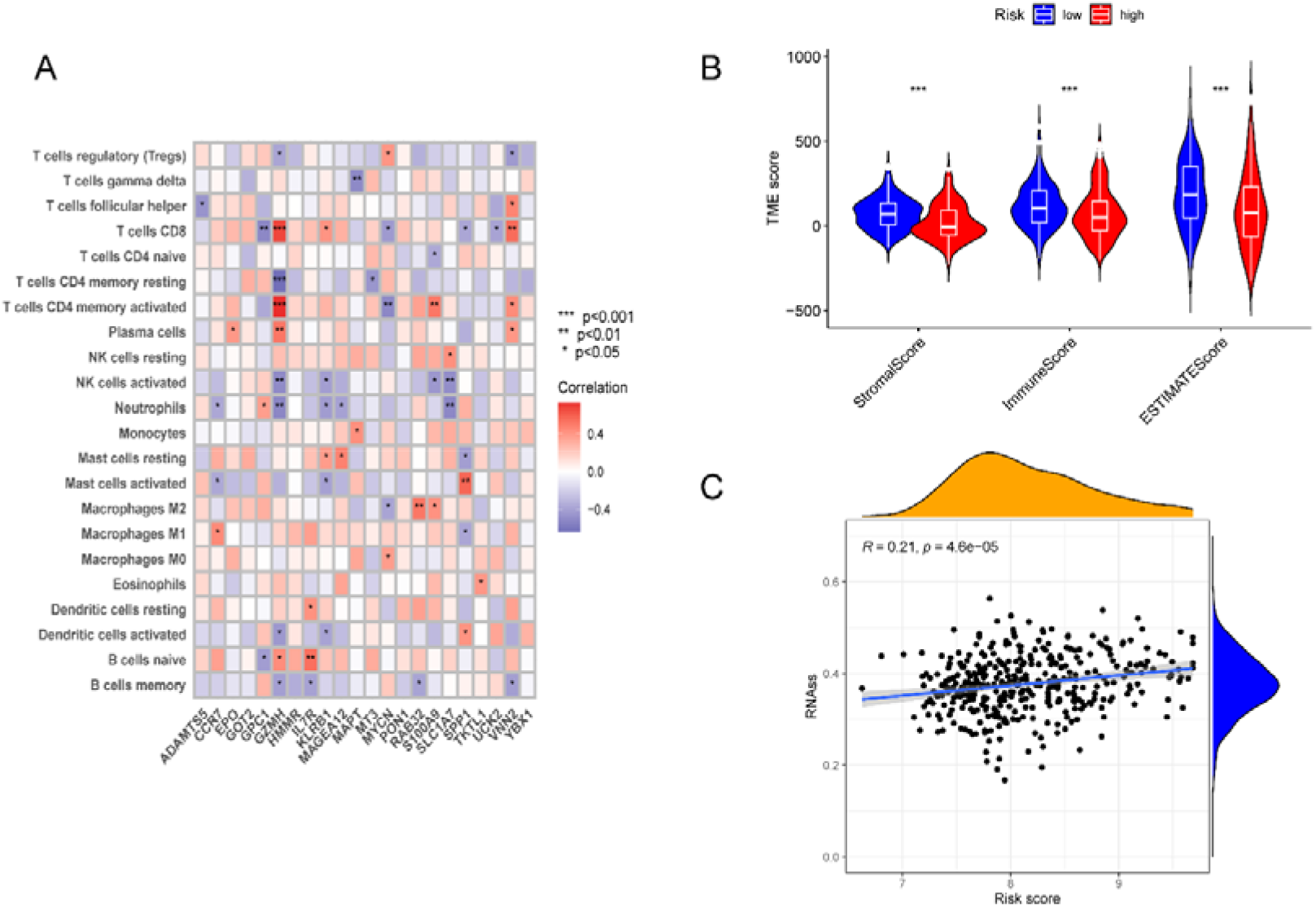
Comprehensive analysis with multiple omics. **A)** The connections between immune cell infiltration and the model genes. The strength of the color represents the degree of correlation. **(B)** The status of the tumor microenvironment in the low- and high-risk groups. **(C)** The correlation of the tumor stem cell index (RNAss) and the risk score (p<0.05).

**Fig. 8:**
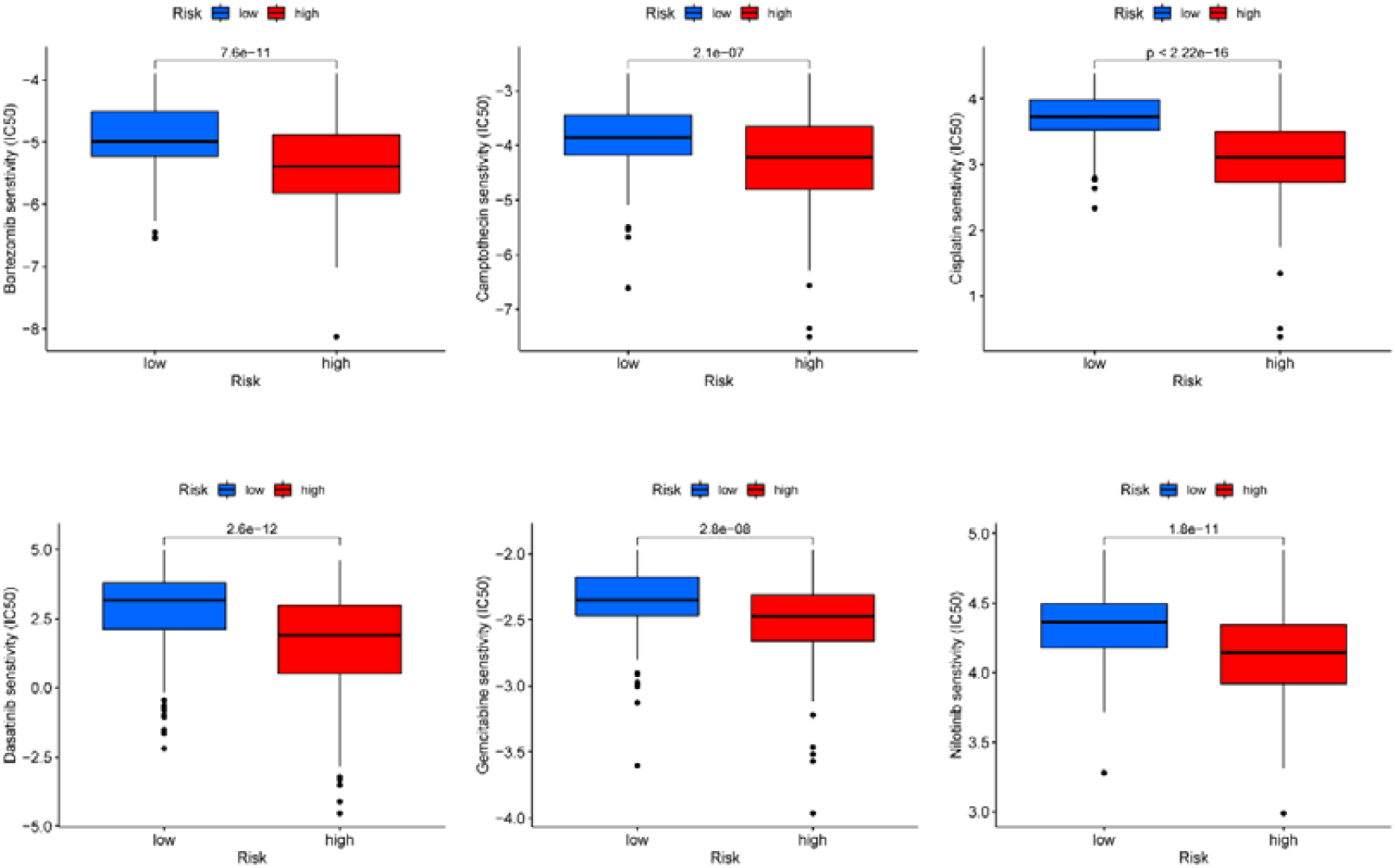
Potential drugs screened by drug sensitivity analysis.

## Discussion

After developing chronic fibrotic liver disease caused by viral or metabolic etiologies, patients tend to develop HCC[13]. However, the key question is how we can reliably estimate the HCC risk and diagnose the early stage of HCC. Unfortunately, a robust estimate system has not been established, and many patients suffer from HCC[14]. To establish a precise prognostic model that can effectively solve this urgent need, we integrated multiomics analysis and the progression of pyroptosis. Pyroptosis is a novel form of cell death and was discovered to have pivotal roles in oncogenesis, immune cell infiltration and antitumor responses[15]. In this study, we collected 52 pyroptosis-related genes and found that most of them have different expression levels between HCC samples and matched normal tissues. This indicated that pyroptosis has important functions in HCC. Furthermore, we explored the protein–protein interactions and mutual regulatory relations of the 42 DEGs. *GSDMD, CASP8, PYCARD* and some CHMP family genes were the core interaction genes. Most of the interacting genes positively regulated each other. Regarding TMB and CNV, *TP53* was the most mutated gene, and the main pattern was missense mutation. We also found that most of the DEGs had CNVs in HCC samples, and *GSDMC, AIM2, GADMD, and CHMP6* had high gain variations, while *CASP9* had high loss variations. Fifty-two prognostic genes were identified using Cox analysis, and the HCC patients could be divided into two clusters using consensus clustering analysis based on the expression level of prognostic genes. It is noteworthy that the two clusters had significant differences in both the OS rate and the clinical features. This differs from other clusters in preparing models for HCC, which have no significant differences in clinical features[16, 17]. This indicates that our method for building the model has more accuracy and significance in predicting the prognosis of HCC patients. Moreover, we integrated multiple omics analysis to demonstrate the landscape of our gene signature and prognostic model in various molecular layers.

Subsequently, we performed LASSO and Cox analyses based on prognostic genes and established a 22-gene signature prognostic model. As expected, this model can separate the different risk patients well, and the two patient subtypes divided by the model had significant OS rate differences. Moreover, the AOCs of ROC analysis for several years are approximately 0.8. Together, these results confirmed that our model is reliable and valuable for predicting HCC patient prognosis. Importantly, the model was well validated in the internal test and external validation cohorts. Furthermore, the univariate and multivariate Cox regression analyses all showed that the risk score generated by the model can serve as an independent prognostic factor. To explore the potential biological functions and pathways that contribute to the risk of developing HCC, we performed GO and KEGG enrichment analyses. We found that material metabolism and molecule secretion could be the mechanism for developing HCC. Recently, Zheng et al. depicted the landscape of tumor-infiltrating T cells across cancers, revealing the heterogeneity of T cells in cancer[18]. Thus, we further evaluated the relevance of infiltrated immune cells and the model genes. *GZMH* was found to have a strong correlation with CD4 and CD8 T cells, which could provide new insight for further study. Regarding the TME, the low-risk group defined by our model had a higher immune-related score, indicating that low-risk patients could benefit from a better immune state[19]. We also clarified that the index of tumor stem cells increased with increasing risk. This result demonstrates that the content of tumor stem cells could be a risk factor for HCC and in turn verifies the robustness of our model[20]. Finally, we screened some drugs that have different sensitivities in treating the two risk groups. The IC50 values of the screened drugs were significantly lower in the high-risk group, illustrating that these drugs have higher sensitivity in the high-risk group than in the low-risk group.

In conclusion, this was the first study to comprehensively investigate the role of pyroptosis in HCC with multiple omics analysis. We established a robust and acute prognostic model for HCC. Compared with other published models, our model showed distinct advantages in multiple aspects. We are the first to integrate the multiomics analysis to establish a pyroptosis-related model. We collected more pyroptosis genes and identified more prognostic genes for building the model. The subtypes showed significant differences in clinical profiles before and after the model was built. All the findings in our study provide a comprehensive landscape of molecular heterogeneity in HCC based on pyroptosis and facilitate the precise management of HCC patients.

## Acknowledgments

National Natural Science Foundation of China (81603119) and Natural Science Foundation of Beijing Municipality (7174316). Major science and technology projects (2018ZX09303047).

## Author Contributions

JHH carried out the experiments, analyzed the data, drew the pictures and wrote the manuscript. YDW and MC conceptualized and designed this study. All authors have revised and agreed to publish this manuscript.

